# Revisiting the gastrin-releasing peptide/bombesin system: A reverse-evolutionary study considering *Xenopus*

**DOI:** 10.1101/2020.05.13.093955

**Authors:** Asuka Hirooka, Mayuko Hamada, Daiki Fujiyama, Keiko Takanami, Yasuhisa Kobayashi, Takumi Oti, Tatsuya Sakamoto, Hirotaka Sakamoto

## Abstract

Gastrin-releasing peptide (GRP), first isolated from the porcine stomach, is a neuropeptide that modulates the autonomic system in mammals and has previously been considered to be the mammalian equivalent of bombesin, a fourteen amino acid peptide first isolated from the skin of the European fire-bellied toad, *Bombina bombina*. Bombesin-like peptides and the related neuromedin B (NMB) have since been identified in mammals. However, the orthologous relationships among GRP/NMB/bombesin and their receptors in vertebrates are still not well understood. Our studies have focused on the GRP system that is widely conserved among vertebrates. We have used phylogenetic analysis and reverse transcription-PCR, quantitative PCR, immunohistochemistry, and Western blotting experiments to examine the expression of both GRP and its receptor (GRPR) in a clawed frog (*Xenopus tropicalis*) and to understand the derivation of GRP system in the ancestor of mammals. We demonstrate, by phylogenetic and synteny analyses, that GRP is not a mammalian counterpart of bombesin and also that, whereas the GRP system is widely conserved among vertebrates, the NMB/bombesin system has diversified in certain lineages, in particular in frog species. In *Xenopus*, we found the expression of the mRNA for both *GRP* and *GRPR* in the brain and stomach. In addition, our quantitative PCR analysis shows that, in *Xenopus*, the expression of *GRP* mRNA is highest in the brain, whereas expression of *GRPR* mRNA is highest in the spinal cord. Our immunohistochemical analysis shows that GRP-immunoreactive cell bodies and fibers are distributed in several telencephalic, diencephalic, and rhombencephalic regions and spinal cord of *Xenopus*. Our Western blotting analysis also indicates the presence of GRPR protein in the brain and spinal cord of *Xenopus*. We conclude that GRP peptides and their receptors have evolved to play multiple roles in both the gut and brain of amphibians as one of the *‘gut-brain peptide’* systems.

**Author Summary:** Bombesin is a putative antibacterial peptide isolated from the skin of the frog, *Bombina bombina*. Two related (bombesin-like) peptides, gastrin-releasing peptide (GRP) and neuromedin B (NMB) have been found in mammals. The history of GRP/bombesin discovery has caused little attention to be paid to the evolutionary relationship of GRP/bombesin and their receptors in vertebrates. We have classified the peptides and their receptors from the phylogenetic viewpoint using a newly established genetic database and bioinformatics. We demonstrate, by phylogenetic and synteny analyses, that GRP is not a mammalian counterpart of bombesin and also that, whereas the GRP system is widely conserved among vertebrates, the NMB/bombesin system has diversified in certain lineages, in particular in frogs. Gene expression analyses combined with immunohistochemistry and Western blotting experiments indicate that GRP peptides and their receptors have evolved from ancestral (GRP) homologues to play multiple roles in both the gut and the brain as one of the *‘gut-brain peptide’* systems of vertebrates, which is distinct from the frog bombesin lineage.

## Introduction

The fourteen-amino acid peptide, bombesin, was initially described as a possible antibacterial peptide isolated from the skin of the European fire-bellied toad, *Bombina bombina*, and was shown to have potent bioactivity in the mammalian nervous system [1, 2]. Subsequently, the mammalian bombesin-like peptides, gastrin-releasing peptide (GRP) [2] and neuromedin B (NMB) [3], were isolated. GRP is a 27-amino acid peptide (29-amino acids in rodents) originally isolated from the porcine stomach as the mammalian equivalent of bombesin [2]. Many studies have indicated that GRP is widely expressed in the central nervous system (CNS) in addition to the gastrointestinal tract in mammals [4, 5]. Because GRP could reproduce most of the biological effects of bombesin in many mammals, GRP had long been considered as the mammalian equivalent of amphibian bombesin [5-7]. Bombesin-like peptides appear to function *via* a family of three G protein-coupled receptors (GPCRs) [8], namely the GRP-preferring receptor (GRPR or BB2 receptor) [9], the NMB-preferring receptor (NMBR or BB1 receptor) [10], and the potential orphan receptor, bombesin receptor subtype-3 (BRS-3 or BB3 receptor) in mammals [11]. To date, in the mammalian CNS, it has been reported that the GRP system might be integral, through GRPR-mediated mechanisms [9], in a variety of autonomic-related functions, including food intake [12-14], circadian rhythms [15-17], fear memory consolidation [18-20], male reproductive function [21], control of sighing [22], and itch sensation [23, 24].

In birds, the expression of orthologous genes for GRP and GRPR has been reported in the chicken CNS [25, 26]. To our knowledge, the only report of a central role of GRP in avian behavior is decreased feeding after the intracerebroventricular administration of GRP in chickens [27]. In amphibians, however, little information is currently available on the function of the GRP/GRPR system in the CNS, because GRP has long been considered as the mammalian equivalent of bombesin [5-7]. Thus, the central role of GRP in non-mammalian species remains unclear. However, it has been reported that frogs synthesize both GRP and bombesin, which are genetically distinct peptides, suggesting that GRP is not mammalian bombesin. In addition, molecular cloning analyses revealed that 3 classes of receptor subtypes were identified in the frog brain [28]. Based on amino acid sequence, two of the classes were clearly the amphibian orthologous genes of the GRPR and NMBR, but not the BRS-3 [28]. Moreover, a fourth class (BB4) of receptor from *Bombina* was identified in amphibians and, interestingly, this had a higher binding affinity for bombesin than for either GRP or NMB [28].

As described above, understanding of the orthologous relationships among GRP/NMB/bombesin and their receptors in vertebrates is still in its infancy. In this study, we classified the peptides and their receptors from the viewpoint of phylogeny by using a newly established genetic database and the bioinformatics approach. In particular, we focus on the GRP system that is widely conserved among vertebrates, and examined its expression in a clawed frog (*Xenopus tropicalis*) to understand its derivation in the ancestor of mammals.

## Results

### Sequence and structure of *GRP* and *GRPR* in *Xenopus*

To verify sequences of *GRP* and *GRPR* in *X. tropicalis*, we cloned cDNA encoding *GRP* and the *GRPR* in *X. tropicalis* and confirmed they are identical to the sequences registered in the GenBank (GRP: XM_018090834.1, GRPR: XP_002938295.1). The deduced amino acid sequence of *Xenopus* prepro-GRP reveals three major components: a signal peptide (31aa in *Xenopus*; 31aa in medaka fish; 27aa in chicks; 23aa in rats; 23aa in humans) (S1 Fig, highlighted in gray); this is followed by the bioactive GRP_1−29_ (mature GRP, GRP_1–24_ in medaka fish; GRP_1–27_ in chicks; GRP_1–29_ in rats; GRP_1–27_ in humans) (S1 Fig, highlighted in pink), including a motif encoding a 10-amino acid-peptide called neuromedin C (NMC or GRP-10; GRP_15–24_ in medaka fish; GRP_20–29_ in chicks; GRP_20–29_ in rats; GRP_18–27_ in humans) at the carboxyl-terminus of mature GRP (S1 Fig, magenta box). The motif is highly conserved in vertebrates; and finally a carboxyl-terminal extension peptide termed pro-GRP_33–121_ in *Xenopus* (pro-GRP_28–99_ in medaka fish; pro-GRP_31–124_ in chicks; pro-GRP_33–124_ in rats; pro-GRP_31–125_ in humans) (S1 Fig, black bar). The mature GRP in *Xenopus* shared high similarity, particularly at the identical [Ser^2^] form-NMC (GRP-10) region, with that in rodents and birds but not with the [Asn^2^] form-NMC in humans and fishes (S1 Fig). Peptide sequences of NMB are highly conserved among vertebrates, particularly 11 amino acids in the carboxyl-terminus are identical between human, rat, chick, medaka, and zebrafish (S1 Fig). Bombesin-like peptides have previously been identified in some frogs; *Rana catesbeiana, R. pipiens* (ranatensin) [29], *Alytes maurus* (alytesin) [30], *Phyllomedusa sauvagii* ([Leu^8^] form-phyllolitorin and [Phe^8^] form-phyllolitorin) [31], *Bombina orientalis* ([Ser^3^, Arg^9^,Phe^13^] form-bombesin, [Phe^13^] form-bombesin, [Leu^13^] form-bombesin) [32, 33] and *Bombina variegate* ([Phe^13^] form-bombesin, [Phe^13^] form-bombesin-like peptide, bombesin, and [His^6^] form-bombesin) [34] (S1 Fig 1, highlighted in blue). Sequences of carboxyl-terminus regions of GPR, NBM and bombesin peptides are quite similar, 4 amino acids (W, A, G, M) are common between these peptides. On the other hands, those of signal peptide region and carboxyl-terminal extension peptide region were diversified between GRP, NMB, and bombesin, and also between the animals.

**Fig 1.**
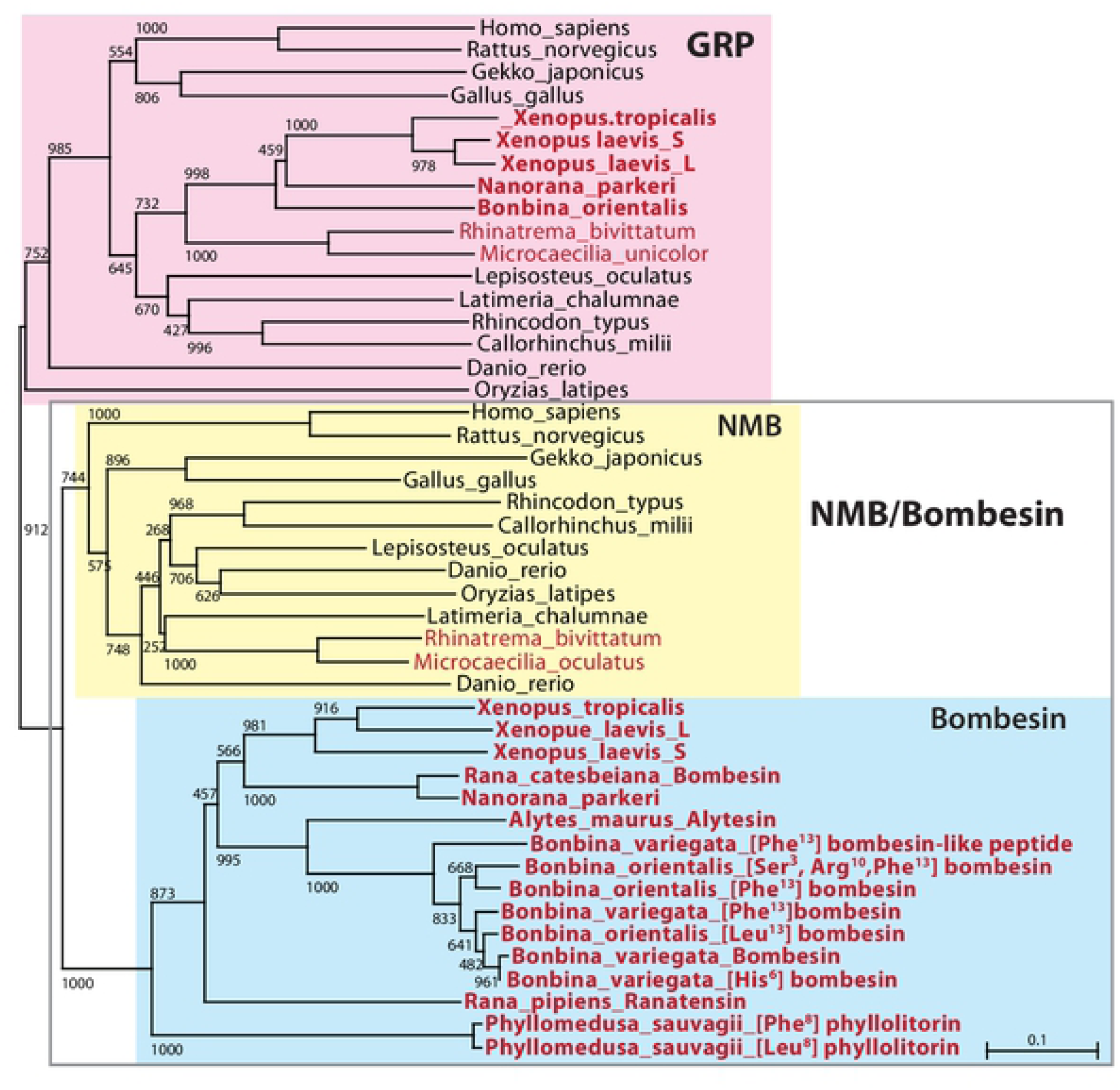
Molecular phylogeny of precursors of GRP/NMB/bombesin family peptides. GRP/NMB/bombesin family peptides are classified into a GRP clade (pink box) and an NMB/bombesin clade (gray frame). While GRP was found in all the animals examined, genes included in NMB clade were not found in frogs (yellow box). All bombesin-like peptides in frogs appeared to form a single bombesin clade (blue box). Species name in red: amphibians; bold: frogs. IDs for the protein sequences used in this analysis are shown in S1Table.

In terms of receptors for bombesin-like peptides, hydrophobicity analysis showed that *Xenopus* GRPR, NMBR, and BRS-3 contain seven transmembrane domains (TM1–7) as well as those in other vertebrates (*e.g.* human, rat, chicken, and zebrafish) (S2 Fig). Homology was highest in the hydrophobic domains, and the [Asp^85^] in the second hydrophobic domain that has possibly been linked to ligand binding is well conserved in vertebrates [35] (S2 Fig).

### Phylogenetic analysis of GRP/NMB/bombesin and GRPR/NMBR/BRS-3

To clarify the relation among GRP/NMB/bombesin family, phylogenetic analysis of GRP/NMB/bombesin precursors (preprohormone) was performed (Fig 1). Homologous genes for GRP/NMB/bombesin family peptides were found in Gnathostomata (higher animals than Cyclostomata/Agnatha)/(vertebrates except Cyclostomata/Agnatha). For the analysis, we used the deduced amino acid sequence in Mammalia *Homo sapiens* and *Rattus norvegicus*, Aves *Gallus gallus*, Reptilia *Gekko japonicus*, Amphibia *X. laevis, X. tropicalis, Nanorana parkeri* (frogs), *Rhinatrema bivittatum* and *Microcaecilia unicolor* (Caecilians), teleost fish *Danio rerio* and *Oryzias latipes*, cartilaginous fish *Callorhinchus milii* and *Rhincodon typus*, because their genomes have been decoded. In addition, the sequences of prepro-bombesin-like peptides, which have been reported for other frog species were also included (Fig 1).

The prepro-GRP/NMB/bombesin sequences were divided into two major clades: GRP and NMB/bombesin clades (Fig 1). A single GRP gene was found in almost all animals examined, although *X. laevis* which is allotetraploid has one gene on each of chromosome L and S (Fig 1, pink box). The NMB/bombesin clade was further divided into the sub-clades: the NMB clade (Fig 1, yellow box) and the bombesin clade (Fig 1, blue box); the bombesin clade and NMB clade were found only in frogs; and all the other animals including caecilians, respectively. *X. tropicalis* and *Nanorana parkeri* have single bombesin gene, and *X. laevis* has the gene on each of chromosome L and S, but these frogs do not possess any genes of the NMB group (Fig 1). In addition, all the precursors of bombesin-like peptides which have previously been identified in other frogs: one in *Rana catesbeiana, R. pipiens*, and *Alytes maurus*; two in *Phyllomedusa sauvagii*; three in *Bombina orientalis*; and four in *Bombina variegate* [34] were also included in the bombesin clade (Fig 1).

These results indicate two possibilities for the evolution of NMB/bombesin: one is the specialization of NMB into bombesin in the frog lineage; the other is the divergence into NMB and bombesin clades resulting, respectively, in the undetectability of frog NMB and the disappearance of bombesin in vertebrates other than frogs. We therefore examined synteny in genes surrounding the amphibian NMB/bombesin locus (Fig 2). Comparison of the genome of *X. tropicalis, Nanorana parkeri, Microcaecilia unicolor* and *Rhinatrema bivittatum*, indicates that the order of genes around the caecilian NMB genes and the frog bombesin genes were highly conserved, although an inversion of the *ZNF11 - SEC592A – bombesin - KTI12* region of *X. tropicalis* genome has occurred. Thus, it can be concluded that bombesin and NMB are respective orthologs and that specialization of the NMB sequence in the frog lineage resulted in bombesin (Fig 2).

**Fig 2.**
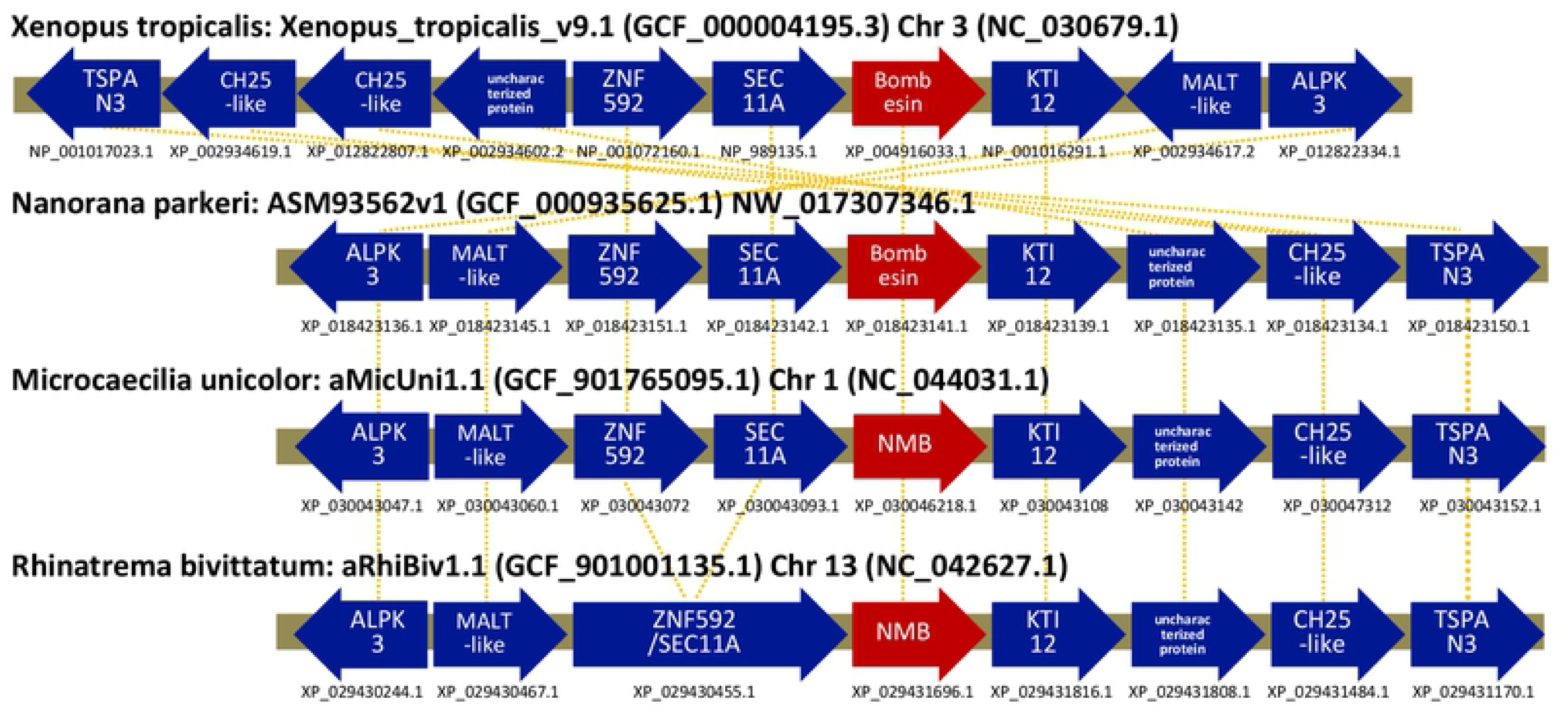
Schematic representation of gene synteny around the NMB/bombesin gene in amphibians. Horizontal lines (gray) indicate chromosome fragments. Genes are represented as arrows (Red: NMB/bombesin gene, Blue: Other genes) according to their transcriptional orientation. Protein IDs are shown below the genes. Yellow lines indicate correspondences of orthologous genes. Gene synteny around the NMB/bombesin gene in amphibians are highly conserved, suggesting an orthologous relationship between NMB in caecilians and bombesin in frogs.

We also performed phylogenetic analysis of the receptors for the GRP/NMB/bombesin family; GRPR, NMBR and BRS-3 (Fig 3). The results suggest that Gnathostomata basically have orthologs in each of the three groups with no specialization in the frog lineage as seen in NMB/bombesin. The BB4 receptor in *Bombina* also belongs to the mammalian BRS-3 group. In addition, the conservation of GRPR and NMB, BRS3 was not found in teleosts (*e.g. Danio rerio* and *Oryzias latipes*) and cartilaginous fish (*e.g. Callorhinchus milii* and *Rhincodon typus*), but only in archaic fish such as *Latimeria chalumnae* (coelacanth), *Erpetoichthys calabaricus* (reedfish), *Lepisosteus oculatus* (gar), and *Acipenser ruthenus* (sturgeon) (S1 Table).

**Fig 3.**
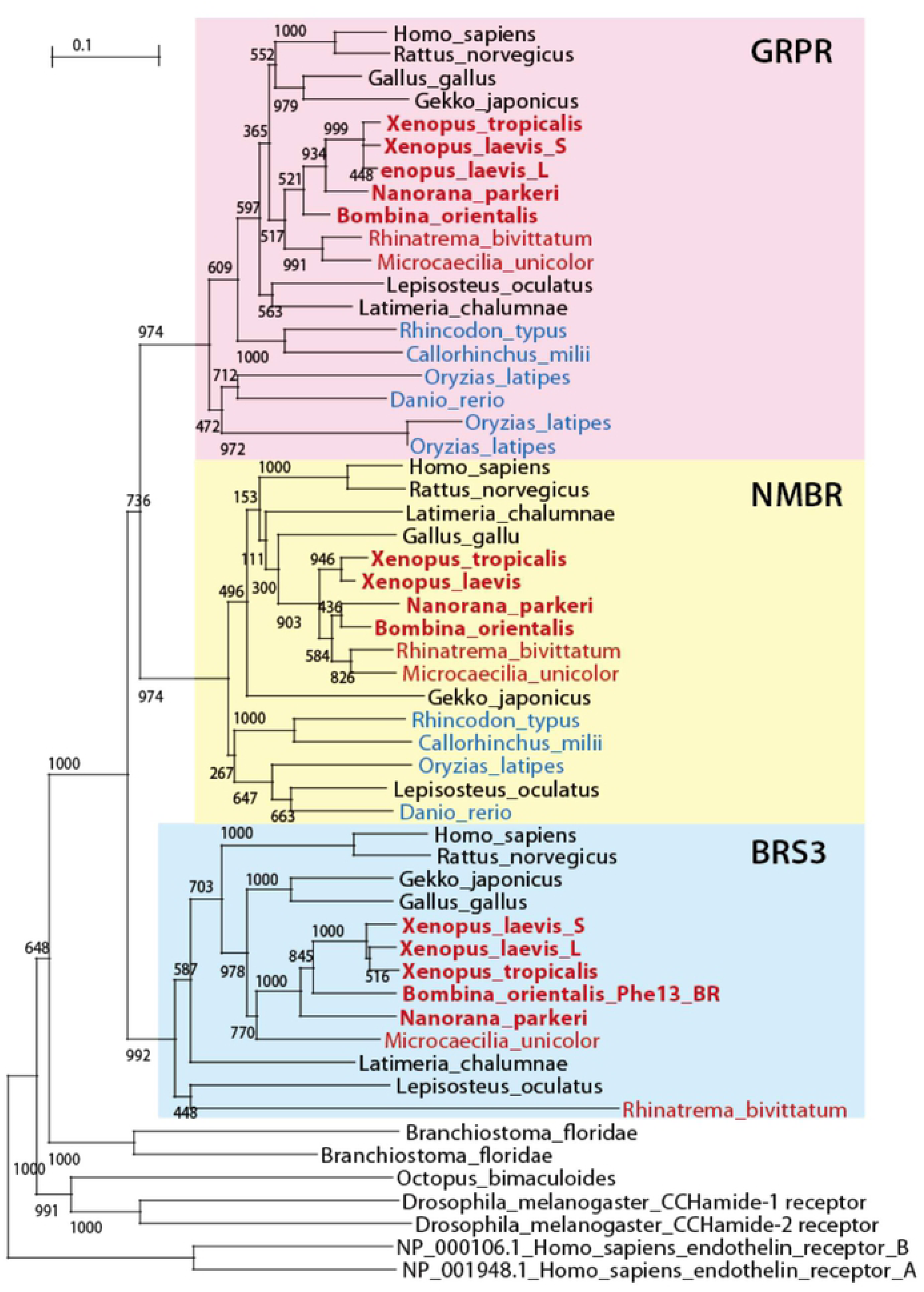
Molecular phylogeny of GRPR/NMBR/BRS-3. **The** GRPR/NMBR/BRS-3 family gene diverged into GRPR (pink box), NMBR (yellow box) and BRS-3 (blue box) in the ancestor of vertebrates. All the three receptors are widely conserved among vertebrates including frogs, although BRS-3 genes were not found in teleost and cartilaginous fish (blue characters). Species name in red: amphibians; bold: frogs. IDs for the protein sequences used in this analysis are shown in S1 Table.

### Reverse transcription (RT)-PCR of *GRP* and *GRPR* mRNA in *Xenopus*

In contrast to the diversification of NMB/bombesin in the frog lineage and the loss of the BRS-3 gene in some fish lineages, GPR and the GRPR are widely conserved in vertebrates. In this study, we used frogs to investigate the principal (conserved, original) role of these bombesin-family systems. We confirmed the expression of *GRP* and *GRPR* mRNA in a variety of *Xenopus* tissues (brain, spinal cord, heart, lung, and stomach) by RT-PCR. Bands were detected at the expected sizes for *GRP* and *GRPR* genes in the brain (Fig 4). *GRP* mRNA was highly expressed in the brain, spinal cord, stomach, and weakly expressed in the lung (Fig 4; *upper panel*). Although *GRPR* mRNA was detected in all tissues, the expression level was low in the heart (Fig 4; *middle panel*). As the internal control in *Xenopus*, nearly equivalent amounts of *GAPDH* cDNA were amplified from RNA preparations among these tissues, which showed that no significant RNA degradation had occurred and a proper RT was obtained (Fig 4; *bottom panel*).

**Fig 4.**
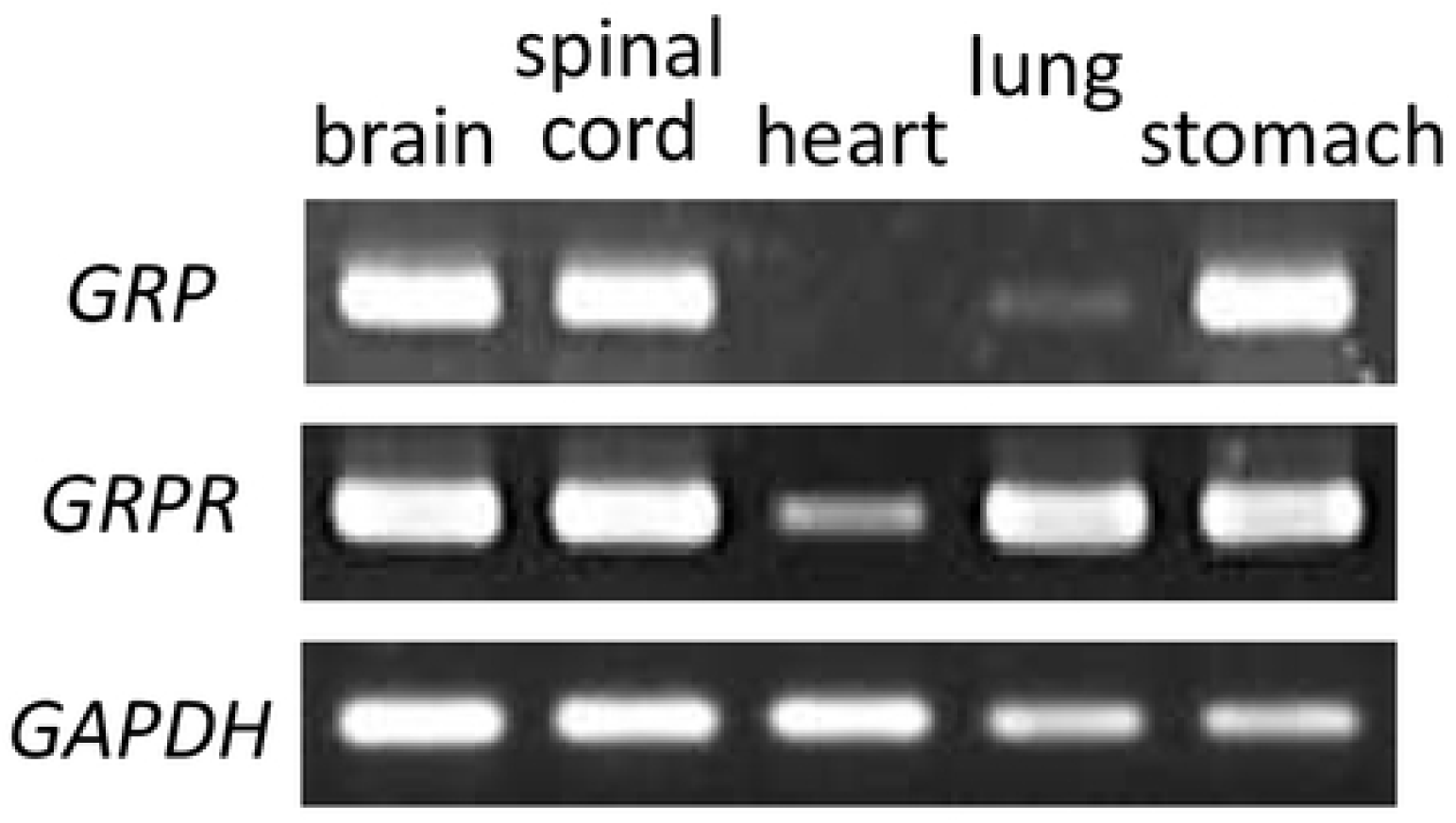
Reverse transcription (RT)-PCR analysis of gastrin-releasing peptide (*GRP*) and GRP receptor (*GRPR*) mRNA expression in *Xenopus tropicalis*. RT-PCR for glyceraldehyde-3-phosphate dehydrogenase (*GAPDH*) was performed as the internal control.

### Real-time quantitative PCR (qPCR) of *GRP* and *GRPR* mRNA in *Xenopus* CNS

To quantify the *GRP* and *GRPR* expression at the transcription level, we performed real-time qPCR analyses for four parts of the central nervous system of males and females: (1) the telencephalon; (2) the diencephalon/mesencephalon/pons/cerebellum; (3) the medulla oblongata; and (4) the spinal cord. Although *GRP* and *GRPR* mRNA expression was detectable in these all tissues of both sexes, no sex differences were detected in any of the tissues we examined (blue bars indicate means of males, red bars indicate means of females) (Fig 5). In both sexes, the expression of *GRP* mRNA was higher in the diencephalon/mesencephalon/pons/cerebellum (2) and the medulla oblongata (3), than in the telencephalon (1) and the spinal cord (4) (Fig 5a). In contrast, the expression of *GRPR* mRNA was the highest in the spinal cord (4), and the lowest in the telencephalon (1) (Fig 5b).

**Fig 5.**
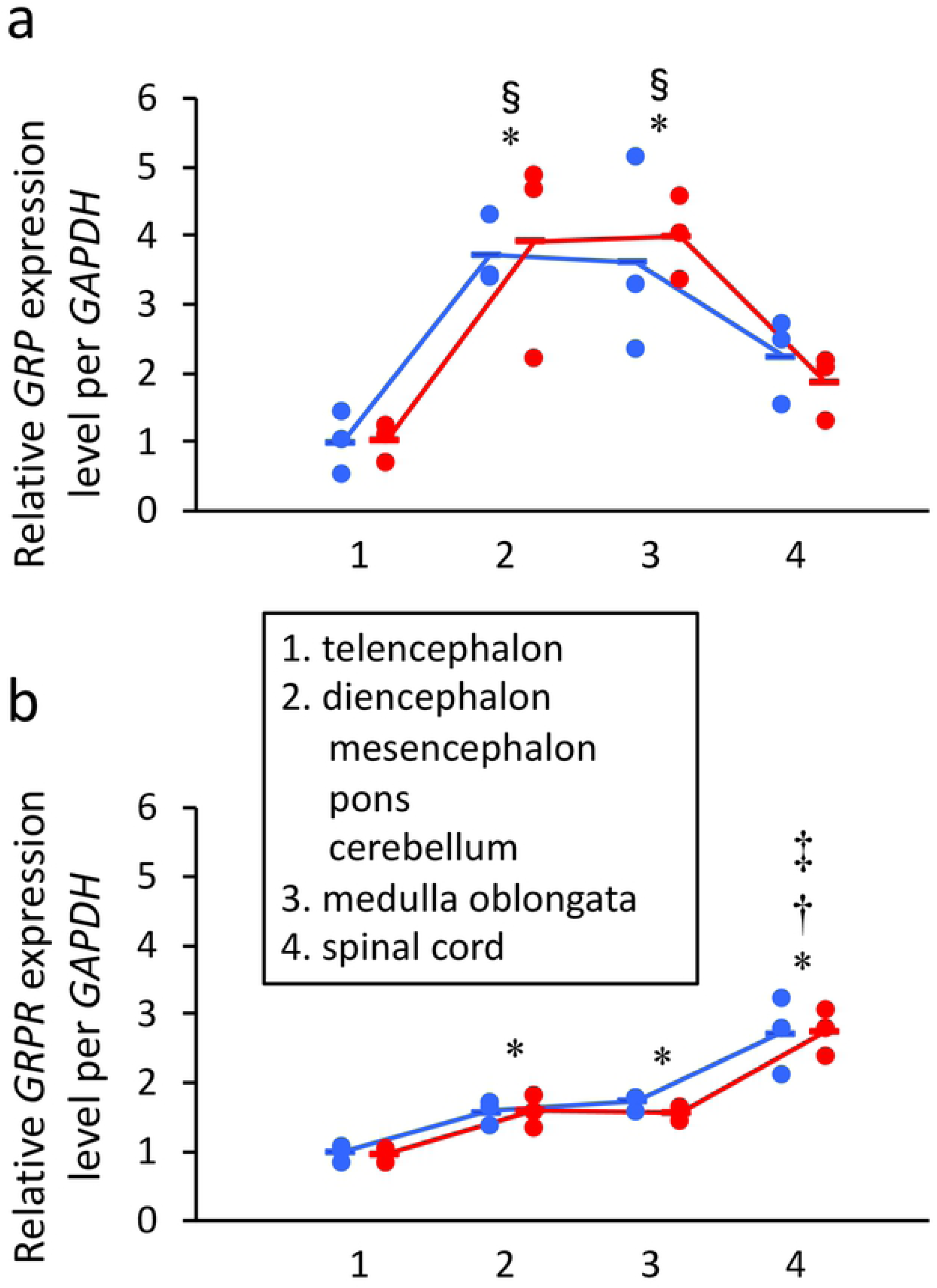
The expression levels of GRP and GRPR mRNA in *Xenopus* central nervous system were measured by the real-time quantitative PCR. Relative expression levels of *GRP* (a) and *GRPR* (b) in the central nervous system: (1) the telencephalon, (2) the diencephalon/mesencephalon/pons/cerebellum, (3) the medulla oblongata, and (4) the spinal cord of males and females were analyzed. P values indicate two-way ANOVA with Tukey’s honestly significant difference test (*vs.* telencephalon (1), \**P* < 0.05; *vs.* diencephalon/mesencephalon/pons/cerebellum (2), †*P* < 0.05; *vs.* medulla oblongata (3), ‡*P* < 0.05; *vs.* spinal cord (4), §*P* < 0.05). No sex differences were detected in any of the tissues. Dots and bars indicate values of each sample and means of the samples, respectively. Blue: males, red: females.

### Distribution of GRP in *Xenopus* CNS

The expression of GRP was next localized in *Xenopus* CNS. The examination of transverse (from rostral to caudal) brain and spinal cord sections revealed the presence of many cell bodies and fibers of GRP-immunoreactive (^+^) neurons in *Xenopus* CNS. The overall neuroanatomical distribution of GRP^+^ neuronal cell bodies and their fiber projections is schematically summarized in Fig 6 and Table 1. The specificity of the GRP antiserum reactivity was confirmed by control absorption experiments in which the primary rabbit antiserum against *Xenopus* GRP_20–29_ was preabsorbed with an excess of *Xenopus* GRP_20-29_ antigen peptide (^2^[Ser] form-NMC); in these experiments no immunostaining was seen (Fig 7). In the spinal cord, GRP^+^ fibers and numerous varicosities were found throughout the spinal grey matter area (Fig 7a, b, d, and e). A cluster of GRP^+^ cell bodies was located mainly in the dorsal field of spinal grey in the cervical spinal cord (df; Fig 7a, b, and e). In the thoracic and lumbosacral spinal cord, similar GRP^+^ fibers were frequently observed, but few GRP^+^ cell bodies could be detected.

**Table 1.**
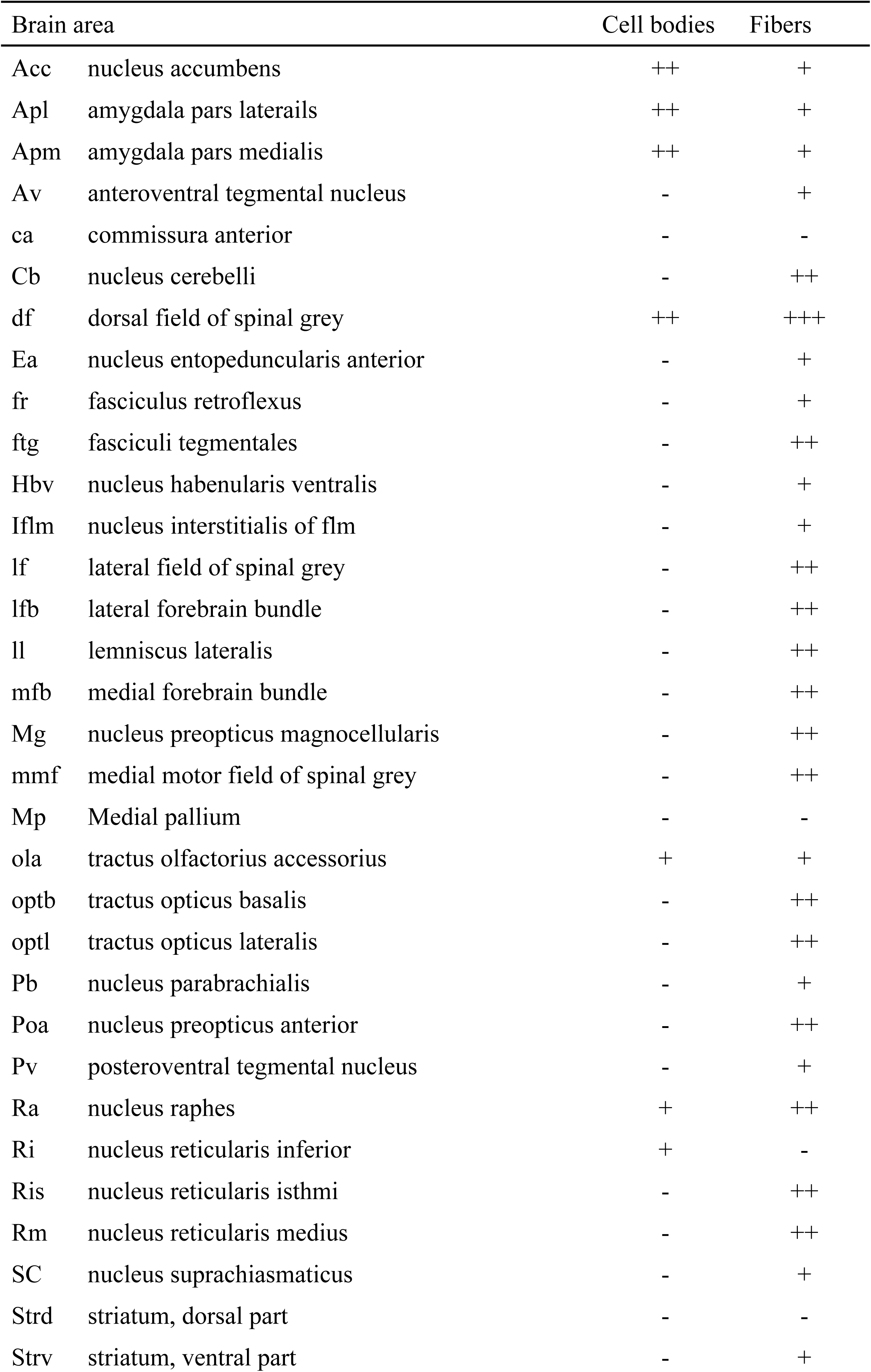
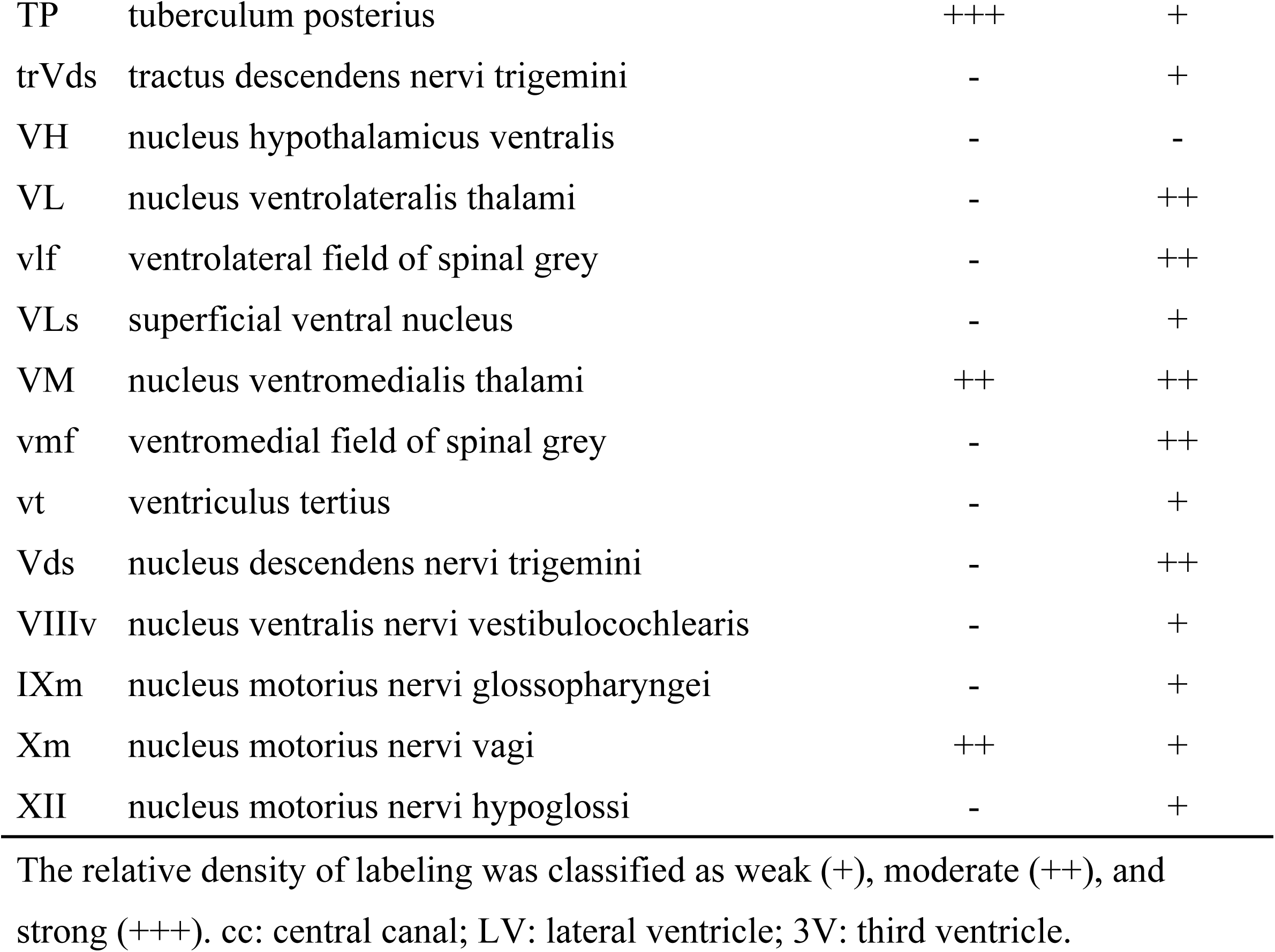
Localization of gastrin-releasing peptide (GRP)-immunoreactive cell bodies and fibers in *Xenopus* central nervous system.

**Fig 6.**
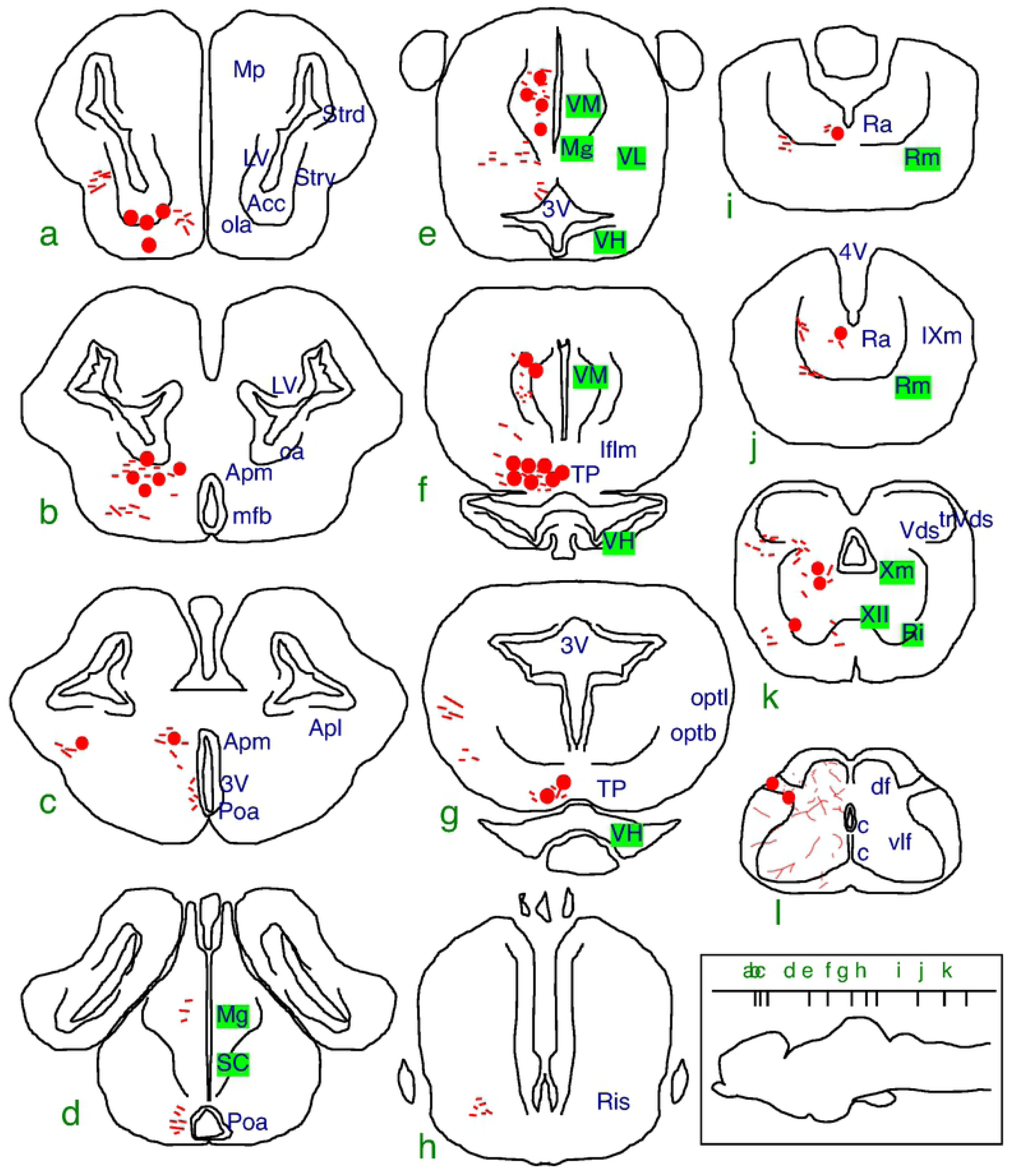
Schematic cross sections illustrating the distribution of GRP-like immunoreactivity in the central nervous system (CNS) of *Xenopus*. A series of illustration of the CNS from telencephalon (a) to cervical spinal cord (l). Immunoreactive cell bodies and fibers in the CNS are shown by red circles and red lines, respectively. The anatomical structures are indicated on the right hemispheres according to the nomenclature of ten Donkelaar HJ [73]. The density of the symbols shown on the left hemispheres is roughly proportional to the relative density of the immunoreactive elements. The letters (a–l) in the schematic lateral profile of the brain (inset) indicates the rostrocaudal level of each transverse section. For abbreviations, see Table 1.

**Fig 7.**
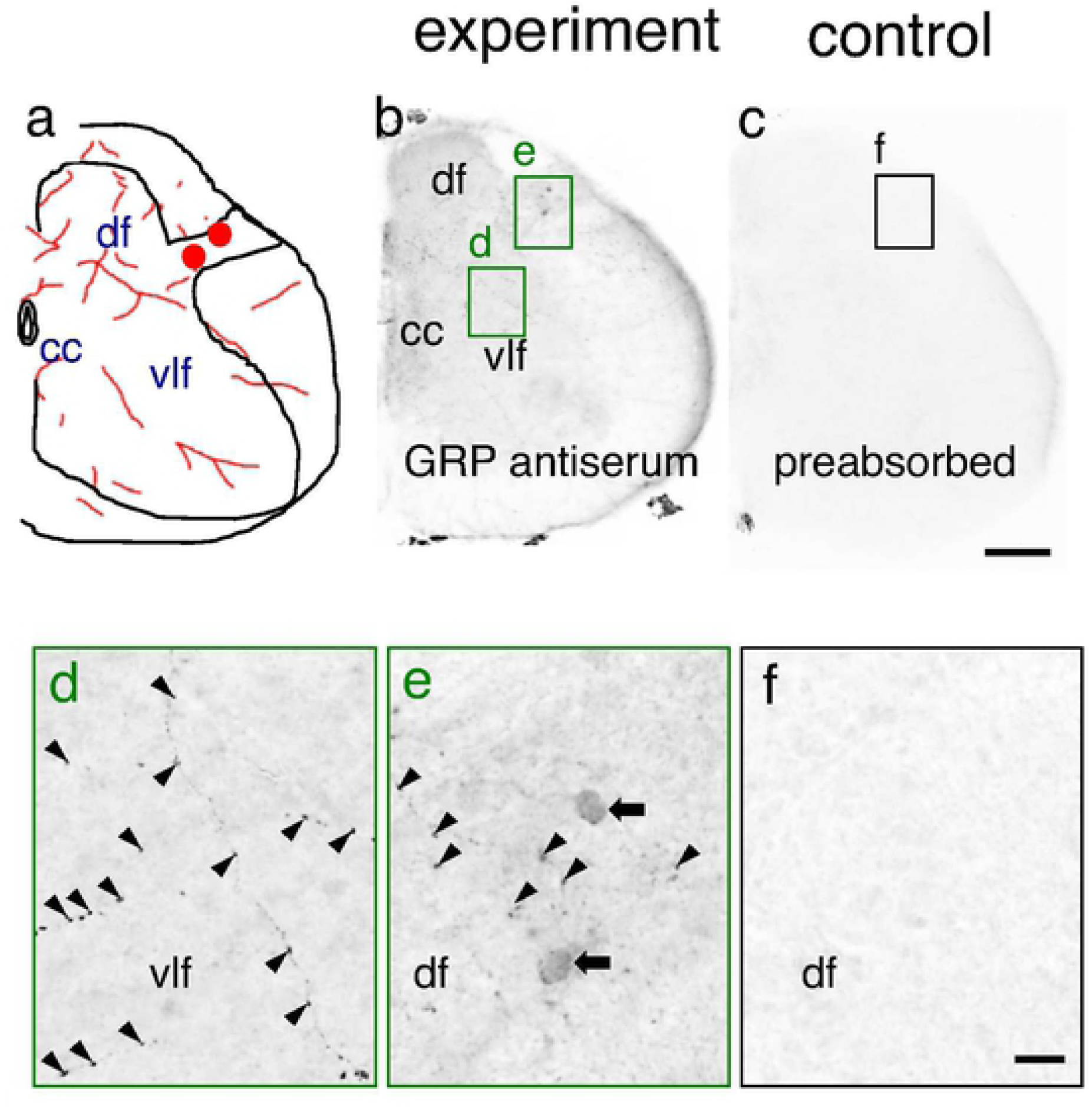
Immunohistochemical identification of gastrin-releasing peptide (GRP) in the cervical spinal cords of *Xenopus*. GRP-immunoreactive cell bodies in the spinal cord were present in the dorsal field of spinal grey (df) (arrows). GRP-immunoreactive fibers and series of varicosities were found throughout the spinal grey (arrowheads). Preabsorbtion of the working dilution (1:2,000) of the primary GRP antiserum with a saturating concentration of GRP antigen peptide [neuromedin C; 50 mg/mL] overnight at 4°C before use eliminated the staining. Scale bars = 100 µm, 10 µm in enlarged images. For abbreviations, see Table 1.

In the telencephalon, abundant but weakly GRP-immunoreactive cell bodies and fibers were located throughout the ventral telencephalic area, *e.g.* cell bodies and fibers in the nucleus accumbens (Acc) and thin fibers in the tractus olfactorius accessorius (ola) (Fig 6a and Fig 8a and c). Immunoreactive cell bodies were abundant in the amygdala pars medialis (Apm; Fig 6b and c and Fig 8c and d), and in the amygdala pars lateralis (Apl; Fig 6c and Fig 8e and f). In the diencephalon, a small number of labeled cells were located in the nucleus ventromedialis thalami (VM; Fig. 6e and f and Fig 8g and h). In the hypothalamus, many intensely labeled cell bodies and fibers were found in the tuberculum posterius (TP; Fig 6 f and g and Fig 8i and j). In the brainstem, large but only weakly immunoreactive cell bodies and fibers were found in the nucleus raphes (Ra; Fig 6i and j and Fig 8k and l), the nucleus motorius nervi vagi (Xm; Fig 6k and Fig 8m and n), and the nucleus reticularis inferior (Ri; Fig 6k and Fig 8o and p).

**Fig 8.**
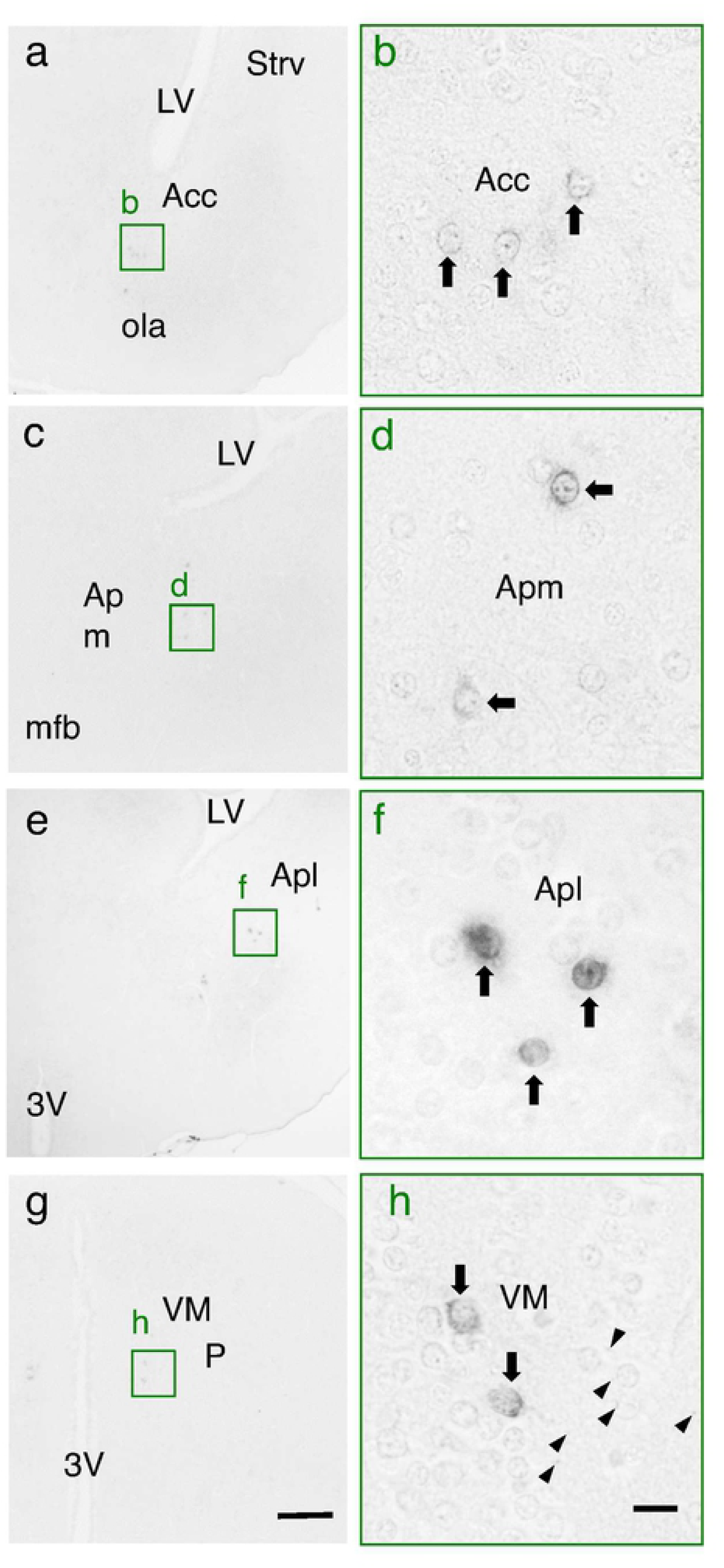
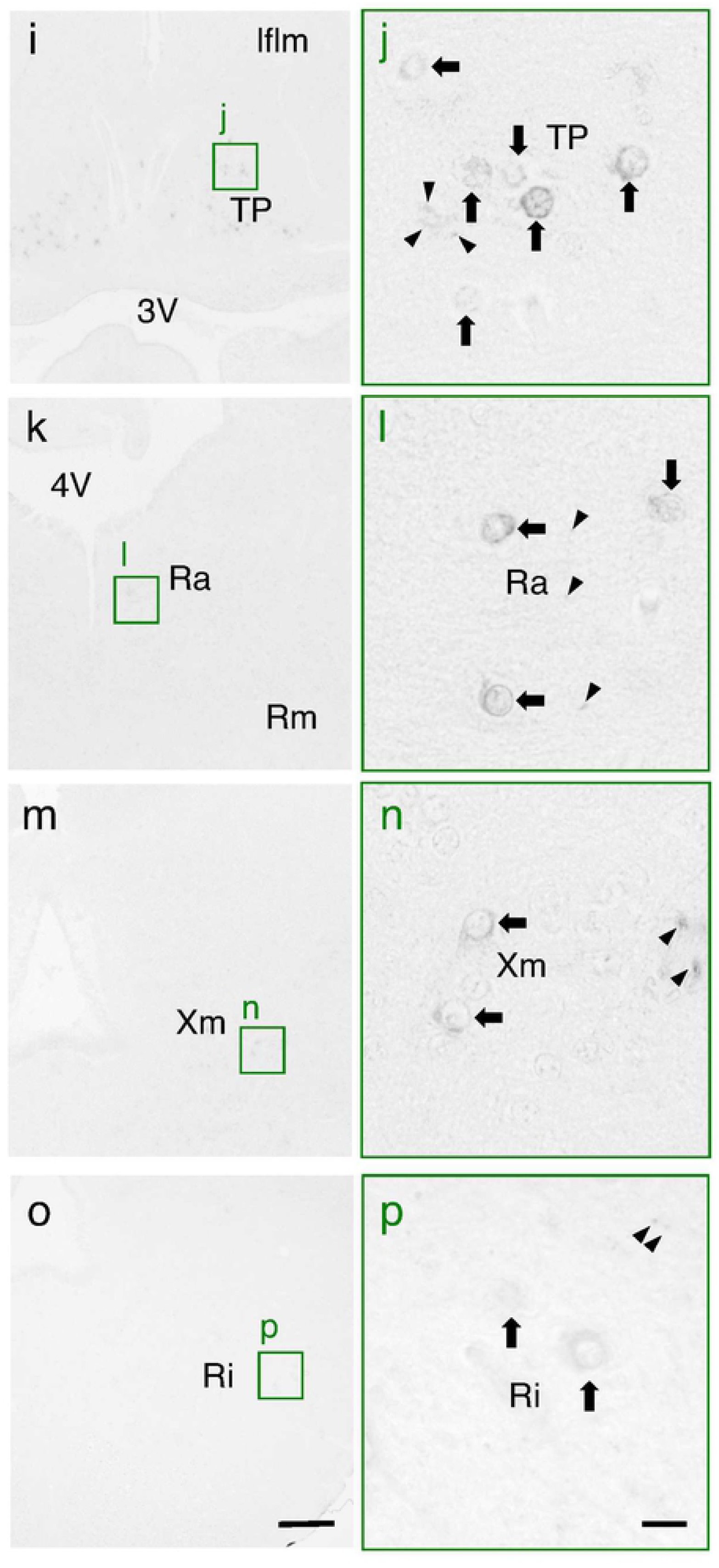
Cell bodies and fibers in *Xenopus* brain immunohistochemically labeled with the anti-gastrin-releasing peptide (GRP) serum. Representative photomicrographs of transverse sections are shown. Dorsal is up and lateral is right. Relative weakly immunoreactive cell bodies and fibers are observed throughout the ventral telencephalic area, *e.g.* in the nucleus accunbens (Acc; a and b). Immunoreactive cell bodies are abundant in the amygdala, *e.g.* in the amygdala pars medialis (Apm; c and d), and in the amygdala pars lateralis (Apl; e and f). In the diencephalon, GRP-immunoreactive somata are detected in the nucleus ventromedialis thalami (VM; g and h) and in the tuberculum posterius (TP; i and j). In the brainstem, GRP-immunoreactive cell bodies and fibers are observed in the nucleus raphes (Ra; k and l), the nucleus motorius nervi vagi (Xm; m and n), and the nucleus reticularis inferior (Ri; o and p). Scale bars = 100 µm, 10 µm in enlarged images. Arrows indicate representative GRP-immunoreactive cell bodies. Arrowheads indicate representative GRP-immunoreactive fibers. For abbreviations, see Table 1.

GRP^+^ fibers were widely distributed throughout the CNS (Fig 6 and Table 1). A fine network of GRP^+^ fibers was also observed in the Apm, in the Apl, and prominently in the striatum, pars ventralis (Strv; Fig 6a and Fig 9a and b). GRP^+^ fibers were also found in some diencephalic nuclei surrounding the third ventricle; *e.g.* VM (Fig 6e and f and Fig 8g and h), nucleus preopticus anterior (Poa; Fig 6c and d and Fig 9c and d), and nucleus preopticus magnocellularis (Mg; Fig 6d and e and Fig 9e and f). A weak distribution of GRP^+^ fibers was also found in the area near the tractus opticus lateralis (optl), the tractus opticus basalis (optb; Fig 6g and Fig 9g and h), and the nucleus reticularis isthmi (Ris; Fig 6h and Fig 9i and j). In the posterior part of the medulla oblongata, the network of GRP^+^ fibers spreading in the nucleus descendens nervi trigemini (Vds) was intensely immunoreactive and such fibers were also scattered in the tractus descendens nervi trigemini (trVds; Fig 6k and Fig 9k and l).

**Fig 9.**
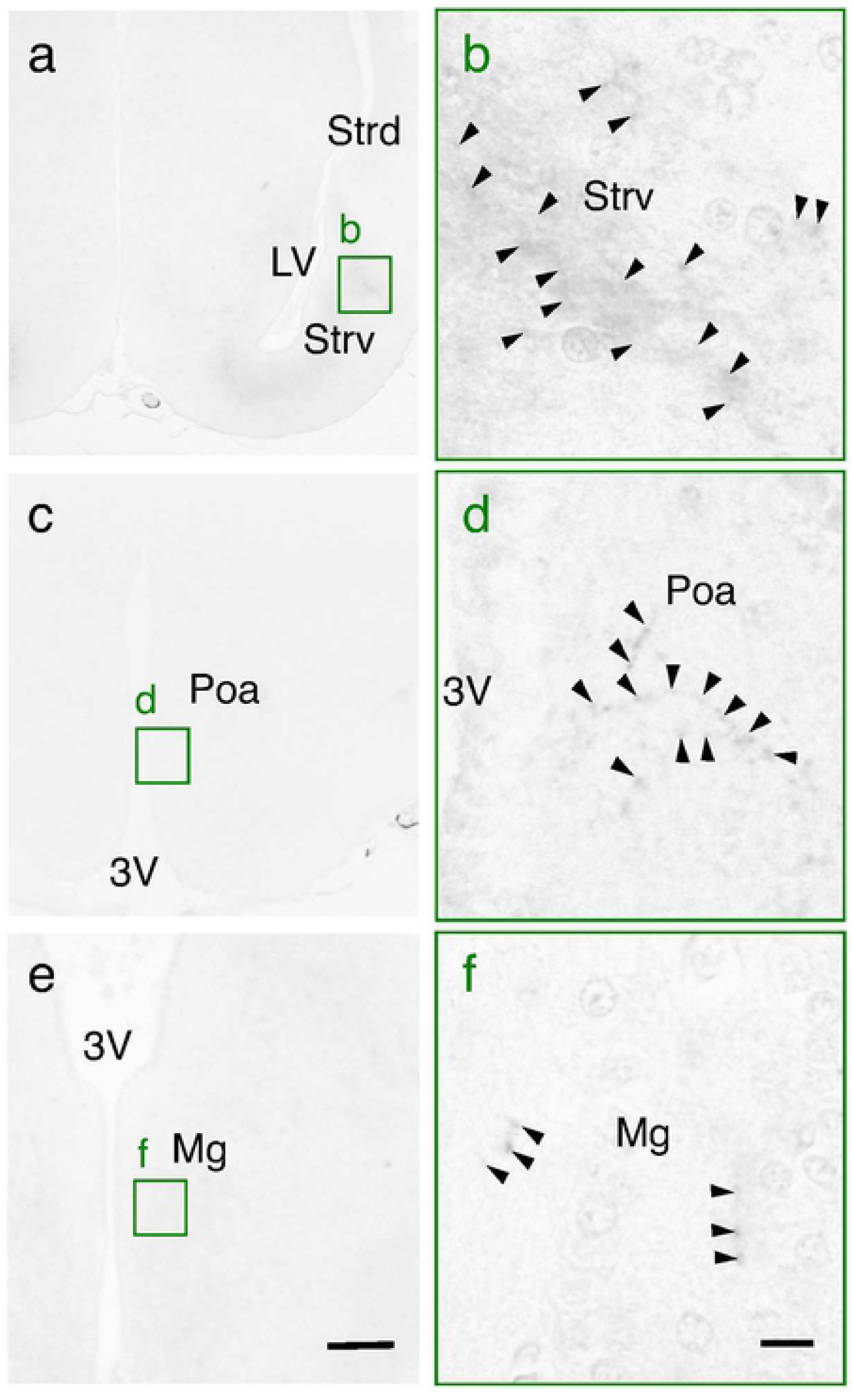
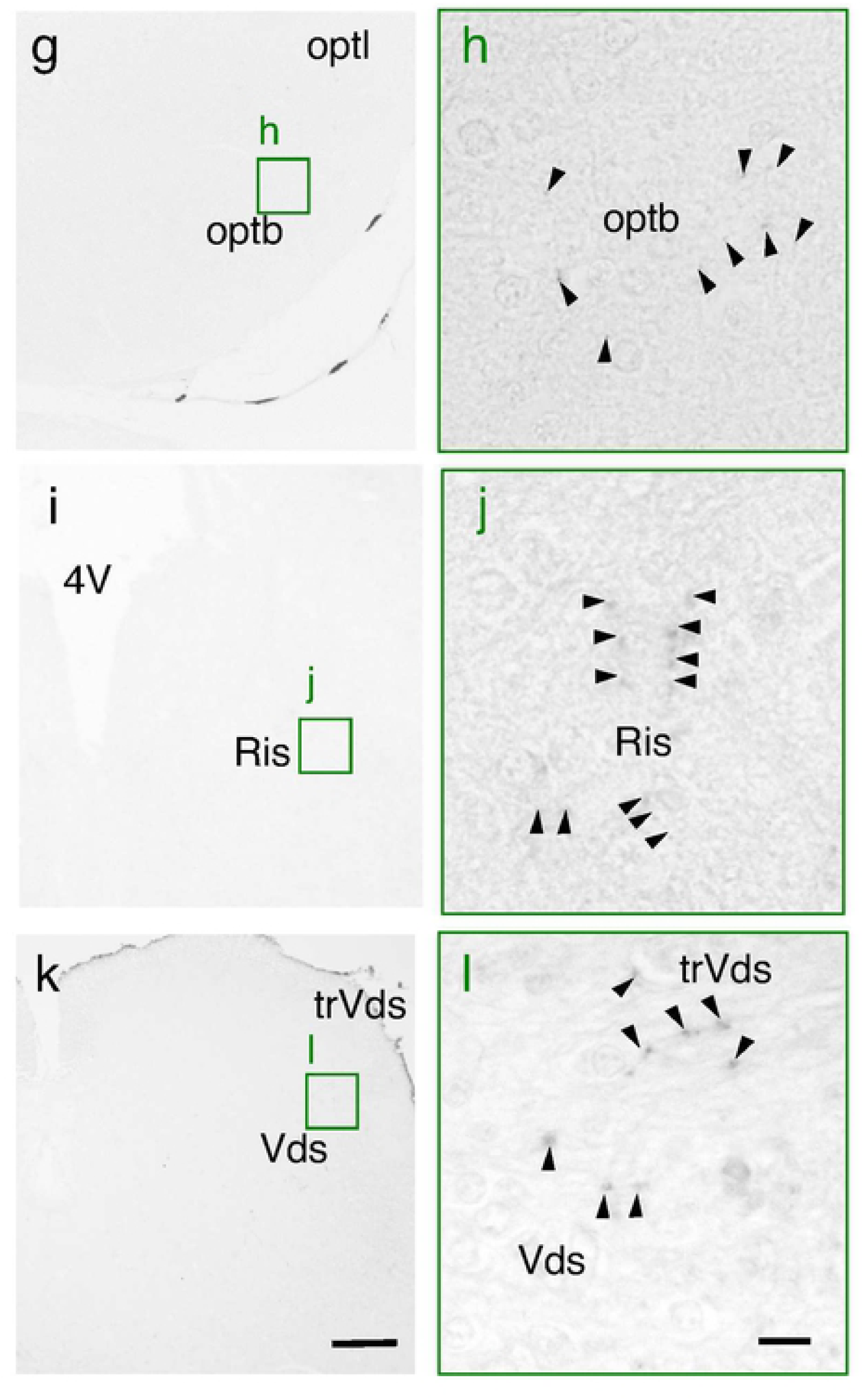
Fibers in *Xenopus* brain immunohistochemically labeled with the anti-gastrin-releasing peptide (GRP) serum. A fine network of GRP-immunoreactive fibers is observed in the striatum, ventral part (Strv; a and b). GRP-immunoreactive fibers are also observed in the preoptic area (Poa; c and d) and in the nucleus preopticus magnocellularis (Mg; e and f). A weak distribution of GRP-immunoreactive fibers is also found in the area near the tractus opticus basalis (optb; g and h), and the nucleus reticularis isthmi (Ris; i and j). In the posterior part of the medulla oblongata, GRP-immunoreactive fibers are observed in the nucleus descendens nervi trigemini (Vds) and in the tractus descendens nervi trigemini (trVds; k and l). Scale bars = 100 µm, 10 µm in enlarged images. Arrowheads indicate representative GRP-immunoreactive fibers. For abbreviations, see Table 1.

### Expression of GRPR protein in *Xenopus* CNS

Western immunoblot analysis with the polyclonal antiserum against *Xenopus* GRPR was performed to determine the presence of GRPR protein in homogenates derived from the brain and spinal cord of adult male *Xenopus*. An intense protein band was observed in the brain and spinal cord extracts, and its electrophoretic mobility was located at ∼43 kDa, which is the expected molecular weight of *Xenopus* GRPR (Fig 10). Preabsorption of the antiserum with an excess of antigen peptides (50 µg/mL) prevented the immunostaining of the ∼43-kDa protein band in the brain and spinal cord (Fig 10). Immunoblot analyses were repeated independently three times by using different three frogs and gave similar results.

**Fig 10.**
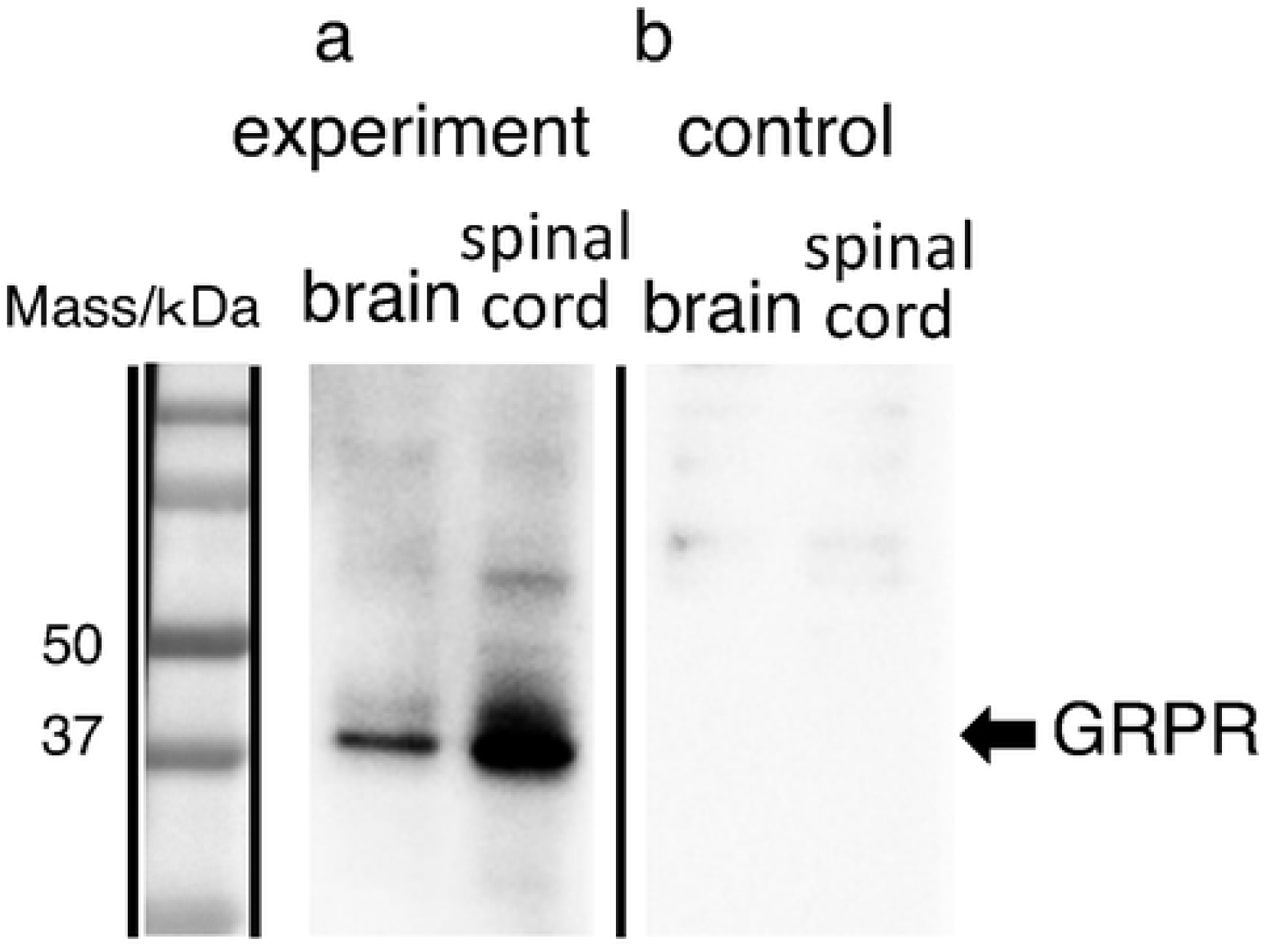
Western immunoblotting of gastrin-releasing peptide receptor (GRPR). The number on the left indicates the molecular weight (kD). Extracts of protein from the *Xenopus* brain and spinal cord were transferred onto polyvinylidene difluoride membranes and probed with the rabbit polyclonal antibody against *Xenopus* GRPR (1:100,000). The antibody recognized a single band at the expected molecular weight of GRPR (∼43 kDa) on a Western blot of the brain and spinal cord. Preabsorption of the antiserum with an excess of antigen peptides (50 µg/ml) eliminated the staining of the ∼43-kDa protein band.

## Discussion

GRP was first identified from the porcine stomach as a mammalian counterpart of the anuran bombesin [2]. Thus, GRP has long been considered as the mammalian equivalents of bombesin [5-7]. It has also been reported that frogs have independent genes for both GRP and bombesin, and this raises the possibility that mammals have an as yet uncharacterized gene encoding a true mammalian bombesin [36]. However, because of the historical background of the GRP discovery [1, 2], little attention has yet been paid to the evolutionary relationship of GRP/bombesin and its receptors in vertebrates. In *Bombina*, it has been suggested that bombesin functions as an antibacterial peptide secreted from the skin [1]. Our phylogenetic analysis indicates that GRP/NMB/bombesin can be divided into two clades; GRP and NMB/bombesin clades (Fig 1). We further found by using synteny analysis that bombesin and NMB are relative orthologs and that specialization of the NMB sequence only in the frog lineage resulted in bombesin (Fig 2). When tetrapods emerged from the sea, they may have started to produce bombesin in the skin as exocrine secretions to protect themselves from bacterial infection and/or predators. To our knowledge, this is the first demonstration of the gene divergence of bombesin and bombesin-like peptides in vertebrates, which appeared to be divided into the two families (*i.e.* GRP and NMB/bombesin) from the vertebrate ancestor (Fig 11).

**Fig 11.**
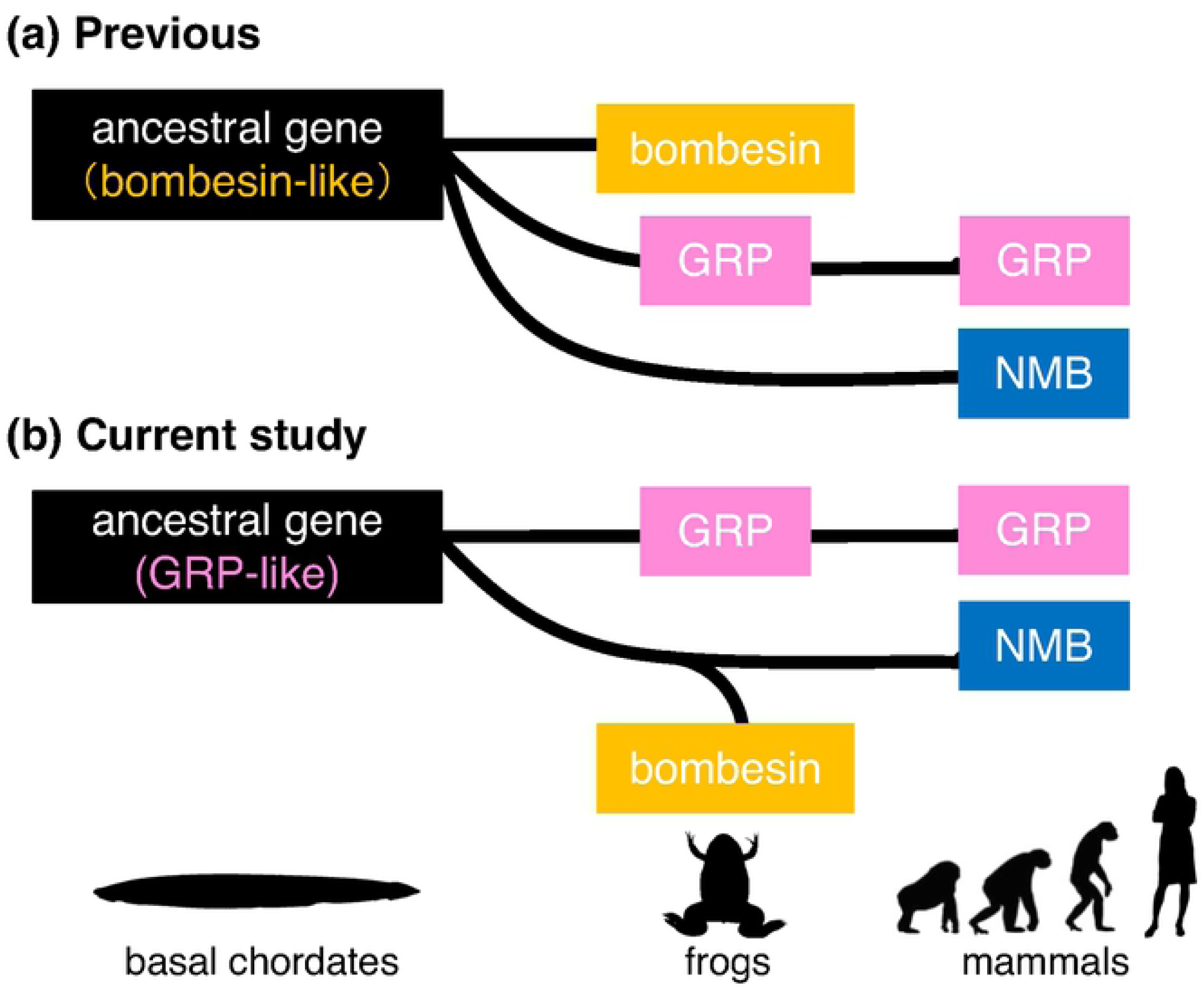
The origin of the gastrin-releasing peptide (GRP)/bombesin lineage. (a) Previous study suggested that GRP (pink box) and neuromedin B (NMB, blue box) should be considered as the mammalian equivalents of bombesin (yellow box). The peptide like bombesin (black box) was thought to be an ancestral form of the bombesin-like family peptides. (b) Our current study demonstrates that GRP is not mammalian counterparts of bombesin and also that, whereas the GRP system is widely conserved among vertebrates, the NMB/bombesin system has diversified in certain lineages, in particular in frog species. The ancestral peptide (black box) is considered to be similar to GRP from the research of the basal chordate amphioxus (Wang et al., 2020).

For bombesin-like peptide receptors, we found that most gnathostomes have an orthologous gene in each of the three groups (*GRPR/NMBR/BRS-3*), although BRS-3 was not found in Teleostei and Chondrichthyes (Fig 3). This suggests that *GRPR/NMBR/BRS-3* diverged into three branches in the ancestor of gnathostomes but that the BRS-3 genes have been lost in Teleostei and Chondrichthyes lineages. Otherwise, it is possible that there were two genes for GRPR and NMBR respectively at the divergence of cartilaginous fish, and that the BRS-3 gene then appeared by duplication of the GRPR gene or NMBR gene in the ancestor of Sarcopterygii while the BRS-3 gene was lost in teleost fish. If this is the case, the amino acid sequence of BRS-3 appears to be specialized, considered from the standpoint of the phylogenetic tree. In addition, no specialization for the receptors was observed in the frog lineage as seen in bombesin-like peptides. In mammals, despite the molecular characterization of BRS-3, determination of its function has been difficult as a result of its low affinity for GRP and NMB and its lack of an identified natural ligand [37-39]. BRS-3-deficient mice develop a mild obesity, associated with hypertension and impairment of glucose metabolism [40]. These results indicate that BRS-3 is required for the regulation of endocrine processes responsible for energy balance in mammals [40]. Because the natural ligand (corresponding to bombesin) for this receptor has never been identified in mammals, birds, or reptiles, BRS-3 is currently considered to be an orphan receptor [37-39]. Thus, BRS-3 might not be essential for life. Knockout mouse studies have also demonstrated that neither *GRPR* nor *NMBR* is on its own essential [9], suggesting that these three bombesin-like peptide receptors can compensate for each others function in mammals. It is often the case that a number of GPCRs with different affinities can be coupled to just one neuropeptide. Recently, it was reported that, in placental mammals, BRS-3 has lost its binding affinity for NMB/GRP and is constitutively active in a ligand-independent manner, in contrast to BRS-3 in non-placental vertebrates including *Xenopus*, which has significant affinity for NMB/GRP [38]. Particularly in *Bombina*, BRS-3 was suggested to be a ‘bombesin-preferring receptor’ [28]. This rather promiscuous relationship between GPCR and ligand could be important in the diversity of cellular functions controlling different life phenomena.

Despite the diversification of NMB/bombesin in the frog lineage and the loss of the BRS-3 gene in some fish lineages, GRP and GRPR are conserved through vertebrates. Peptides synthesized in endocrine cells of the gastrointestinal tract and in neurons are traditionally considered not only as modulators of metabolism, energy balance, appetite, *etc*., but also as neuromodulators (neuropeptides); so called ‘*gut-brain peptides*’ in mammals [41]. GRP appears to be one of gut-brain peptides, and the GRP system might play multiple roles in both the gut and the brain [5]. Indeed, Holmgren *et al*. [42] reported immunohistochemical evidence that the myenteric plexus and circular muscle layer of the stomach of the mudpuppy (*Necturus maculosus*), a salamander, are richly innervated by GRP^+^ fibers, and Kim *et al*. [43] reported that endogenous GRP potently stimulates the contraction of longitudinal muscle and relaxes circular muscle in *Xenopus* stomach, suggesting that GRP is important in the regulation of gastric motility in *Xenopus*. Furthermore, in *Xenopus*, we found expression of the mRNA for both *GRP* and *GRPR* in the brain and stomach, suggesting that *Xenopus* GRP systems play physiological roles locally in the CNS as well as in the stomach. In addition, our qPCR analysis suggests that, in *Xenopus* CNS, the expression of *GRP* mRNA is highest in the brain, whereas *GRPR* mRNA expression was highest in the spinal cord. Our immunohistochemical analysis shows that GRP^+^ cell bodies are distributed in several telencephalic, diencephalic, and rhombencephalic regions and spinal cord of *Xenopus* (see Fig. 6 and Table 1). In particular, we observed many GRP^+^ cell bodies in the hypothalamus and putative limbic system in *Xenopus*, which corresponds well with the mammalian case [5]. GRP^+^ fibers were also distributed widely throughout the CNS of *Xenopus* (Fig 6 and Table 1). Thus, GRP might play an important role in multiple physiological functions in *Xenopus* CNS. In mammals, it is reported that orofacial pruriceptive inputs are processed mainly in the superficial layers of the trigeminal sensory nucleus caudalis in the medulla oblongata, which is similar to the spinal dorsal horn [44]. Therefore, GRP/GRPR signaling in the trigeminal ganglion-trigeminal sensory nuclei of the medulla oblongata appears to play an important role in orofacial itch sensation in mammals [45, 46]. We also found that, in *Xenopus*, abundant GRP^+^ fibers are distributed in the trVds and Vds areas, which appear to correspond to the trigeminal somatosensory system in mammals. Thus, GRP may modulate neurotransmission and integration of somatosensory information in *Xenopus*. Taken together, these results indicate that GRP functions not only as a gastrointestinal bioactive peptide but also as a neuropeptide in *Xenopus* CNS, and that GRP functions as a gut-brain peptide in both amphibians and mammals.

In conclusion, GRP has long been considered as the mammalian equivalents of bombesin [6, 7] (Fig 11a). We now demonstrate, by phylogenetic and synteny analyses, that GRP is not a mammalian counterpart of bombesin, and that the GRP system is widely conserved throughout vertebrates, whereas the NMB/bombesin system diversified in some lineages (Fig 11b). Furthermore, we demonstrate that the GRP system might play multiple roles both in the gut and in the brain of amphibians as one of the ‘*gut-brain peptide*’ systems. Indeed, it has recently been reported that the expression of the common ancestral genes for *GRP/NMB/bombesin* (possibly *GRP* as an ancestral gene) in amphioxus (which belongs to the subphylum Cephalochordata, an extant representative of the most basal chordates) is abundant in the gut, and is also observed in the cerebral vesicle that has been proposed as the homologue of the vertebrate brain [47]. However, the original roles of GRP as an ancestral gene remains unclear at the moment and further functional studies by using lower vertebrates are needed to draw a firm conclusion.

## Materials and Methods

### Ethics statement

All experimental procedures were approved in accordance with the Guide for the Care and Use of Laboratory Animals prepared by Okayama University (Okayama, Japan). All efforts were made to minimize animal suffering and reduce the number of animals used in this study.

### Animals

Male and female adult clawed frogs (*X. tropicalis*), the Golden strain, were provided by the Amphibian Research Center (Hiroshima University, Japan) through the National Bio-Resource Project of the Japan Agency for Medical Research and Development (AMED), Japan. Frogs were maintained according to well established protocols.

### Phylogenetic analysis and gene synteny analysis

For a molecular phylogenetic analysis of precursors of GRP/NMB/bombesin family peptides and GRPR/NMBR/BRS3, protein sequences were obtained from NCBI protein database. Multiple alignments were produced with CLUSTALX (2.1) with gap trimming [48]. Sequences of poor quality that did not well align were deleted using BioEdit [49]. Phylogenetic analyses were performed using the Neighbor-Joining method [50] by CLUSTALX with the default parameters (1,000 bootstrap tests and 111 seeds). Representative phylogenetic trees were drawn by using NJ plot [51]. Signal peptide site in GRP/NMB/bombesin and transmembrane domain in GRPR/NMB/BRS-3 were predicted by SignalP-5.0 [52] and TMHMM2.0 program [53].

For comparison of gene synteny around the NMB/bombesin gene in the amphibian genome, the genome regions upstream and downstream of NMB/bombesin genes were examined using assembled genome sequences of aMicUni1.1 (GCF_901765095.1) for *Microcaecilia unicolor*, aRhiBiv1.1 (GCF_901001135.1) for *Rhinatrema bivittatum*, ASM93562v1 (GCF_000935625.1) for *Nanorana parkeri, X. tropicalis*_v9.1 (GCF_000004195.3) for *X. tropicalis*.

Genome and amino acid sequences of *Homo sapiens, Rattus norvegicus* [54], *Gallus gallus* [55], *Gekko japonicus* [56], *X. laevis* [57], *X. tropicalis* [58], *Nanorana parkeri* [59], *Rhinatrema bivittatum* [60, 61], *Microcaecilia unicolor, Danio rerio* [62], *Oryzias latipes* [63], *Callorhinchus milii* [64], *Rhincodon typus* [65], *Rana catesbeiana, Rana pipiens* [29], *Alytes maurus* [30], *Phyllomedusa sauvagii* [31], *Bombina orientalis* [32, 33], *Bombina variegate* [34], *Latimeria chalumnae* [66, 67], *Erpetoichthys calabaricus* [68], *Lepisosteus oculatus* [69], *Acipenser ruthenus* (Cheng et al., 2019), *Branchiostoma floridae* [70], *Octopus bimaculoides* [71] and *Drosophila melanogaster* [72] used for the analysis were obtained from GenBank. Accession IDs were shown in S1 Table.

### cDNA cloning of *GRP* and *GRPR* in *Xenopus*

Adult male frogs (*n* = 2) were anesthetized with 1% MS-222 (tricaine, Sigma-Aldrich, St. Louis, MO, USA) and sacrificed by decapitation. Immediately, dissected tissues (hypothalamus, spinal cord, and stomach) from frogs were fixed with RNA*Later* solution (Ambion, Austin, TX, USA) and stored at −30°C until RNA extraction. Total RNA was extracted from samples using the illustra RNAspin Mini RNA Isolation kit (GE Healthcare, Buckinghamshire, UK) according to the manufacture’s protocol. The concentration of total RNA was measured using a Qubit RNA assay kit (ThermoFisher Scientific, Waltham, MA, USA). In order to identify genomic sequences of *Xenopus* GRP and GRPR, the 1^st^ cDNA was synthesized from 200 ng of total RNA of stomach origin by using oligo-dT primers and an iScript cDNA Synthesis Kit (Bio-Rad Laboratories, Hercules, CA, USA). The sequences of primers for cDNA cloning were designed based on the genome resource (see Table 2). The resulting RT-PCR amplicons of the stomach sample (full open reading frame sequence for *GRP* or partial sequence for *GRPR*) were subcloned into the pGEM-T easy vector (Promega, Madison, WI) followed by transfection into *Escherichia coli* DH5α competent cells (Takara Bio, Shiga, Japan). Positive clones were identified by blue-white screening and at least three positive clones were sequenced. The primer sequences used in this study are shown in Table 2.

**Table 2.**
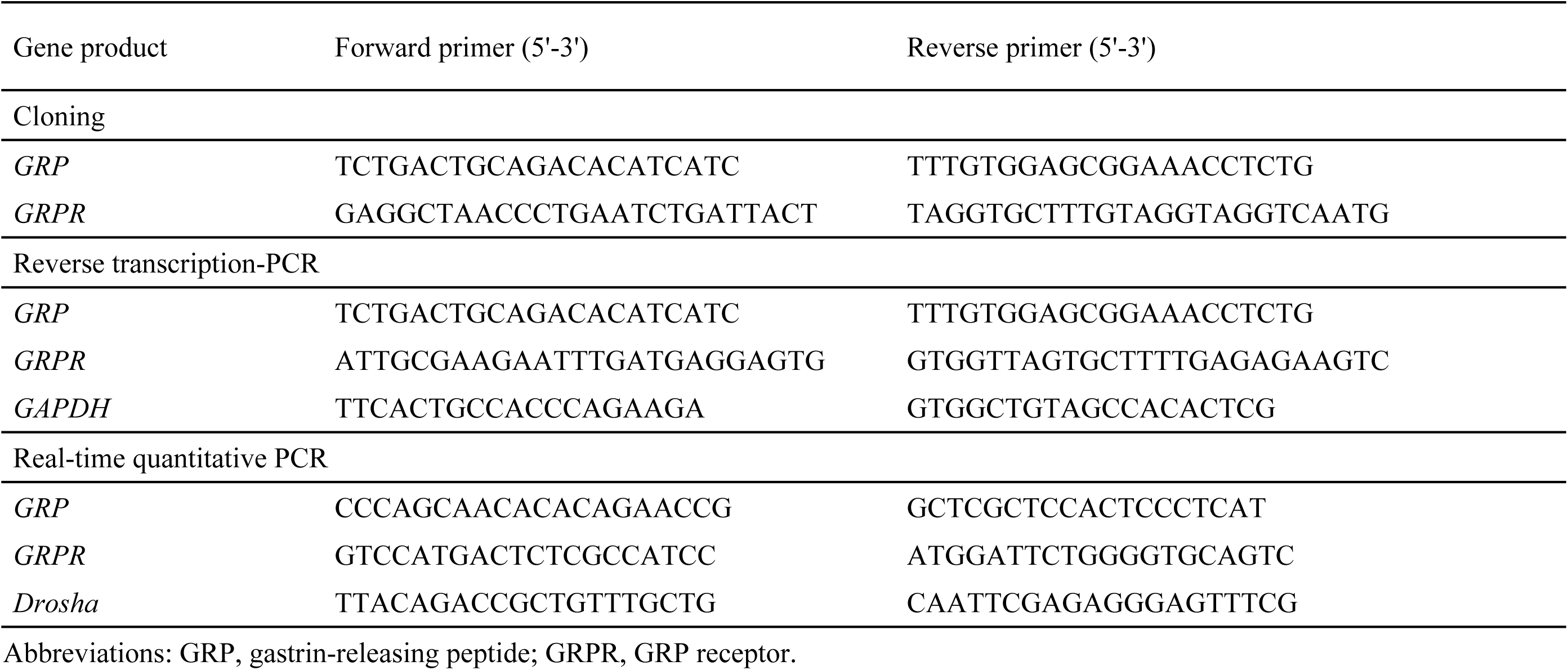
Primer sequences for cloning, reverse transcription-PCR, and real-time quantitative PCR.

### RT-PCR

To determine the tissue distribution of *GRP* and *GRPR* mRNA in *X. tropicalis* (*n* = 3 of each sex), RT-PCR analysis was performed. Total RNA was extracted from various tissues (brain, spinal cord, heart, lung, stomach), using illustra RNAspin Mini RNA Isolation Kit (GE Healthcare). First-strand cDNA was synthesized from 200 ng of total RNA in a 20-µL reaction volume using oligo-dT primers and iScript RT (Bio-Rad Laboratories).

Glyceraldehyde-3-phosphate dehydrogenase (GAPDH) was used as the internal control. Primer pairs are shown in Table 2. The resultant PCR amplicons were electrophoresed on 2% agarose gels. RT-PCR studies were repeated four times using independently extracted RNA samples from different animals. Consistent results were obtained from each run.

### Real-time qPCR

The quantification of *GRP* and *GRPR* in *Xenopus* CNS was performed using real-time qPCR. In brief, whole brains and spinal cords of males or females were quickly removed and placed on ice (*n* = 3 of each sex). Brains were divided into 3 parts: 1 the telencephalon; 2 the diencephalon/mesencephalon/pons/cerebellum; and 3 the medulla oblongata. The preparations were carefully dissected under a dissecting microscope (Olympus, Tokyo, Japan). Total RNA was isolated from samples using illustra RNAspin Mini RNA Isolation Kit with RNase-free DNase I (GE Healthcare) according to the manufacture’s protocol. Samples were reverse transcribed from 500 ng of total RNA in a 20 µL volume using iScript cDNA synthesis kit (Bio-Rad Laboratories). Real-time qPCR was carried out on a C1000™ Thermal Cycler (Bio-Rad Laboratories). Reactions were performed in 20 µL solution, with 200 nM of each primers, 1 µL of 5 ng cDNA samples and SYBER-green master mix (KAPA SYBER FAST qPCR kit, KAPA Bio-systems, Boston, MA, USA) according to the manufacture’s instructions. Assays (in triplicate) were repeated at least twice with the constitutive *Drosha* as a normalizing control. Primer pairs are shown in Table 2. The data relative changes in mRNA expression were determined using the 2^−ΔΔCT^ method. Statistical tests were performed using IBM SPSS ver. 24 for Windows (IBM, Armonk, NY, USA).

### Immunohistochemistry

Adult male frogs (*n* = 4) were killed by decapitation under deep anesthesia (see above). Brains and spinal cords were immediately dissected out and immersion-fixed in Bouin’s fixative solution [saturated picric acid:10% unbuffered formalin:acetic acid = 15:5:1 (v/v)] overnight at 4°C. Subsequently, tissues were dehydrated and embedded in paraffin wax. Serial sections of each tissue were cut transversely or sagittally on a microtome at 10-µm in thickness. We performed immunohistochemical analysis according to our established methods [21]. In brief, endogenous peroxidase activity was eliminated from the sections by incubation in a 0.1% H_2_O_2_ in absolute methanol solution for 20 minutes followed by three 5-minute rinses with phosphate-buffered saline (PBS) (pH 7.4). After blocking nonspecific binding with 1% normal goat serum and 1% BSA in PBS containing 0.3% Triton X-100 for 30 minutes at room temperature, sections were then incubated in Can Get Signal A (TOYOBO) with a 1:2,000 dilution of primary rabbit antiserum against rat GRP_20–29_, a 10-amino acid-peptide called NMC or GRP-10 (AssayPro, St. Charles, MO, USA) for 48 hours at 4°C. This GRP antiserum has previously been shown to be specific for rodent GRP [21, 45]. The amino acid sequence of NMC (epitope residues) is identical between frogs and rodents. Immunoreactive-products were detected with a streptavidin-biotin kit (Nichirei, Tokyo, Japan), followed by diaminobenzidine (DAB) development according to our previous method [21]. GRP-expressing cells in the spinal cord were localized using an Olympus FSX100 microscope. Nomenclature of brain areas and nuclei was based on the stereotaxic atlas of the CNS of anurans [73].

Control procedures for the DAB method were performed using pre-absorption of the working dilution (1:2,000) of the primary antiserum with saturating concentration of frog GRP_20–29_ antigen peptide, (GSHWAVGHLM; 50 µg/mL: ^2^[Ser]-NMC, produced in AnaSpec; San Jose, CA) overnight at 4°C before use. The information of GRP antibody used in this study is showed in Table 3.

**Table 3.** Primary antibodies used in this study.

| Antigen | Description | Source, host species, cat#, or code# | Working dilution | RRID |
| --- | --- | --- | --- | --- |
| GRP | Synthetic peptide mapping at 20–29 amino acids (GSHWAVGHLM) of rat GRP | AssayPro, rabbit polyclonal, 11081-05015 | 1:2,000 | AB_2571636 |
| GRPR | Synthetic peptide mapping at 195–208 amino acid residues (CAPYPHSNGLHPRI) of *Xenopus* GRPR | Generated by our laboratory, rabbit polyclonal, XGR14 | 1:100,000 | AB_2832951 |

Abbreviations: RRID, Research Resource Identifier; GRP, gastrin-releasing peptide; GRPR, GRP-preferring receptor.

### Western immunoblotting

Western blotting was conducted according to our previously described methods [74]. In brief, adult male frogs (*n* = 3) were sacrificed by decapitation under deep anesthesia (see above). Brains and spinal cords were quickly removed and placed on ice, and were snap-frozen immediately in liquid nitrogen. The preparations (100 µg protein/lane) were boiled in 10 µL sample buffer containing 62.5 mM trishydroxymethyl-aminomethane-HCl (Tris-HCl; pH. 6.8), 2% SDS, 25% glycerol, 10% 2-mercaptoethanol, and a small amount of bromophenol blue. Samples were run on a 4–20% SDS-PAGE and electroblotted onto a polyvinylidene difluoride (PVDF) membrane (Bio-Rad Laboratories) using a semidry blotting apparatus (Bio-Rad Laboratories). Membranes were blocked with PVDF Blocking Reagent for Can Get Signal (TOYOBO, Tokyo, Japan) for 30 minutes at room temperature and incubated for 1 hour at room temperature in Can Get Signal Solution 1 (TOYOBO) with a 1:100,000 dilution of rabbit polyclonal antibody against GRPR. The polyclonal antibodies were produced in a rabbit using a *Xenopus* GRPR fragment peptide. The sequence of antigen peptide was 195–208 amino acid residues (CAPYPHSNGLHPRI) of *Xenopus* GRPR (GenBank accession No. XP_002938295.1). Blotted membranes were washed three times with 0.05% Tween 20 in Tris-HCl buffered saline (TBST) and incubated with horseradish peroxidase-conjugated goat polyclonal antibody against rabbit IgG (Bio-Rad Laboratories) at 1:10,000 dilution in Can Get Signal Solution 2 (TOYOBO) for 1 hour at room temperature. After washing for three times with TBST, blots were visualized by Immun-Star WesternC Chemiluminescence Kit (Bio-Rad Laboratories). Images of the different immunoblots were slightly adjusted in brightness and contrast to provide a uniform background. Control procedures were performed by pre-absorption of the working dilution (1:100,000) of the primary antiserum with a saturating concentration of *Xenopus* GRPR_195–208_ antigen peptide, CAPYPHSNGLHPRI (50 µg/mL, produced in Sigma-Aldrich) overnight at 4°C before use. The information of GRPR antibody used in this study is showed in Table 3.

## Acknowledgments

We are grateful to the National Bio-Resource Project (NBRP) of ‘*X. tropicalis*’ in the Amphibian Research Center (Hiroshima University, Japan) for providing *X. tropicalis*, Golden strain. We are grateful to Prof. John F. Morris (University of Oxford, UK) for valuable discussion and for reading this manuscript.

## Author Contributions

A.H., D.F., K.T., and T.O. performed immunohistochemical experiments. A.H. and K.T. performed Western blotting. A.H., M.H., Y.K., and T.O. performed molecular biology, phylogenetic, and gene synteny analyses. T.S. interpreted the data and provided the advice. M.H. and H.S. wrote the paper. A.H. and M.H. contributed equally to this study. H.S. supervised the whole study. All authors had full access to all the data in the study and take responsibility for the integrity of the data and the accuracy of the data analysis.

## Funding

This work was supported by Japan Society for the Promotion of Science (JSPS) KAKENHI (to H.S.; 15K15202, 15KK0257, 15H05724, 16H06280)

(https://www.jsps.go.jp/english/e-grants/index.html), the Japan Agency for Medical Research and Development (AMED) (to H.S.; 961149)

(https://www.amed.go.jp/en/aboutus/index.html), and Okayama University Dispatch Project for Female Faculty members (M.H.)

(https://okayama-u-diversity.jp/grant-support-activities/dispatch-project-female-faculty/). K.T. and T.O. were supported by Research Fellowships of JSPS for Young Scientists (https://www.jsps.go.jp/english/e-pd/index.html).

## Legends for Supplementary Figures

**S1 Fig. Amino acid sequence alignment of gastrin-releasing peptide (GRP), neuromedin B (NMB) and bombesin.** The signal peptide in each sequence by SignalP is shown in a gray box. Bioactive GRP (mature GRP) and neuromedin C (NMC) /GRP-10 site are highlighted in pink and a magenta box respectively. Bioactive NMB in human and rat and bombesin-like peptides previously found in frogs are highlighted in yellow and blue respectively. Carboxyl-terminal extension peptide sites are represented by black line. IDs for the protein sequences used in this analysis are shown in S1 Table.

**S2 Fig. Amino acid sequence alignment of gastrin-releasing peptide-preferring receptor (GRPR), neuromedin B-preferring receptor (NMB) and bombesin receptor subtype-3 (BRS-3).** The seven transmembrane domains (TM1-7) in each sequence identified by TMHMM2.0 program are shaded in gray. IDs for the protein sequences used in this analysis are shown in S1 Table.

